# Perceptual weighting of binaural lateralization cues across frequency bands

**DOI:** 10.1101/2019.12.20.873059

**Authors:** Axel Ahrens, Suyash Narendra Joshi, Bastian Epp

**Affiliations:** Hearing Systems Section, Department of Health Technology, Technical University of Denmark

**Keywords:** Binaural hearing, Spectral integration, Spectral weighting, Auditory enhancement

## Abstract

The auditory system uses interaural time- and level differences (ITD and ILD) as cues to localize and lateralize sounds. The availability of ITDs and ILDs in the auditory system is limited by neural phase-locking and by the head size, respectively. Although the frequency-specific limitations are well known, the relative contribution of ITDs and ILDs in individual frequency bands in broad-band stimuli are unknown. To determine these relative contributions, or spectral weights, listeners were asked to judge the lateralization of stimuli consisting of eleven simultaneously presented 1-ERB-wide noise bands centered between 442 Hz and 5544 Hz and separated by 1-ERB-wide gaps. Interaural disparities were applied to each noise band and were roved independently on every trial. The weights were obtained using a multiple linear regression analysis. In a second experiment the effect of temporal context on the spectral weights was investigated. Ten of the noise bands were used as pre- and postcursors and listeners were asked to lateralize the stimuli. Results show that only the lowest- or highest frequency band received highest weight for ITD and ILD, respectively. Temporal context led to significantly enhanced weights given to the band without the pre- and postcursor. The weight enhancement could only be observed at low frequencies, when determined with ITD cues and for low and high frequencies for ILDs. Hence, the auditory system seems to be able to change the spectral weighting of binaural information depending on the information content.

**PACS:** 43.64.-q *·* 43.64.+r *·* 87.19.lt *·* 43.66.-x *·* 43.66.+y *·* 43.64.Bt

## 1 Introduction

An important ability of the auditory system is spatial hearing. This ability enables the localization of sound sources in auditory space (see Middlebrooks and Green 1991, for a review) and to improve the understanding of speech in environments with interfering sound sources (e.g. Bronkhorst 2000). The two binaural cues available to the auditory system are interaural disparities in time (interaural time difference, ITD) and interaural disparities in level (interaural level difference, ILD). ITDs are predominantly used at low frequencies and ILDs are predominantly used at high frequencies Rayleigh (1907). The spectral dominance of ITD and ILD is commonly referred to as the Duplex Theory and are widely acknowledged in the literature (e.g. Macpherson and Middlebrooks 2002). It is, however, less clear what the contribution of the different cues in the different frequency bands is in realistic conditions with broadband stimuli and with dynamic spectro-temporal context.

At low frequencies the ITD is a reliable cue and small changes in ITD can be detected. At high frequencies, the ITD of the fine structure becomes ambiguous (Rayleigh 1907; Blauert 1984; Moore 2014) and hence unreliable as a cue. For narrow band signals, ITD detection thresholds have been shown to be lowest between 700 and 1000 Hz and to increase towards lower and higher frequencies (Klumpp and Eady 1956; Brughera et al. 2013). The upper frequency limit of ITD detection was shown to be at about 1.5 kHz (Moore 2014; Zwislocki and Feldman 1956; Klumpp and Eady 1956; Brughera et al. 2013). At frequencies above 1.5 kHz, ITD cues in the envelope of a signal can be detected when imposed on carrier frequencies well above 1.5 kHz (Henning 1974; McFadden and Pasanen 1976; Bernstein and Trahiotis 1994; Nuetzel and Hafter 1976; Leakey et al. 1958). At high frequencies, the wavelength of the sound waves becomes small in comparison to the size of the head. This leads to a reduction in sound intensity at the contralateral ear relative to the ipsilateral ear and leads to ILD cues (Blauert 1984). ILD detection thresholds have been shown to be approximately constant over a broad range of frequencies (Yost 1988; Grantham 1984; Rowland and Tobias 1967).

Previous studies have suggested that the frequency-specific detection thresholds of the interaural disparities are related to their relative contribution to spatial hearing. One approach used to derive the spectral weighting is to invert discrimination thresholds for narrowband signals and to calculate sensitivity under the assumption that high sensitivity indicates a high relative contribution, and hence a high weighting for the cue in that frequency band. For ILDs, this approach leads to a constant ILD weighting across frequency bands. For narrow band noises ITD weighting is maximal between 700 and 1000 Hz and decreases towards high and low frequencies (Raatgever 1980; Stern et al. 1988; Buchholz et al. 2018).

However, detection (McFadden and Pasanen 1976) and discrimination (Heller and Richards 2010; Trahiotis and Bernstein 1990) thresholds of ITD and ILD as well as the lateralization extent (Heller and Trahiotis 1996) are affected by the presence of signals in remote spectral regions, a phenomenon known as binaural interference. For example, the ITD threshold of a probe in a frequency band is increased in the presence of an interfering signal in a frequency band lower than the probe frequency (Best et al. 2007). Thus, the spectral weights obtained by inverting thresholds for narrowband signals in isolation might not be applicable for broadband signals.

Furthermore, Buell and Hafter (1991) showed that binaural information are summed across frequency bands if the information belong to the same auditory object. Thus, when multiple auditory objects of target and interfering signal are formed, no binaural interference occurs. In addition, it has been shown that pre- and post-cursors can reduce the detection threshold of a masked signal and make the signal perceptually ‘pop out’, also referred to as auditory enhancement (Viemeister 1980; Byrne et al. 2011). This effect might also affect the spectral weighting of the interaural disparities.

In the present study, we investigated how the auditory system integrates binaural information across frequency for broadband signals. An observer response weighting analysis paradigm was used to determine relative contributions of spectral bands to sound source lateralization. Previously, this analysis has been used to estimate the spectral and temporal weights for the judgement of spectral shape (Lutfi and Jesteadt 2006; Berg 1990), for spectral weights of loudness (Jesteadt et al. 2014; Joshi et al. 2016; Leibold et al. 2007, 2009; Oberfeld et al. 2012), and for temporal weights of ITDs and ILDs (Stecker and Hafter 2002; Brown and Stecker 2010, 2011; Stecker et al. 2013; Stecker 2014; Stecker et al. 2018; Dye et al. 2005). Using an observer-response weighting analysis enables the estimation of weights using stimuli with interaural disparities above threshold and taking binaural interference into consideration. In the first experiment, the spectral weights for ITD and ILD were derived by imposing conflicting ITD and ILD information in multiple frequency bands and asking listeners to judge the lateralization of the stimulus. In a second experiment, the spectral weights of ITDs and ILDs were derived in the presence of pre- and post-cursors to investigate the effect of a dynamic spectro-temporal context.

## 2 Methods

### 2.1 Experiment 1: Spectral weights in static condition

Figure 1 shows a schematic of the stimuli used in experiment 1. Each bar indicates a 1-ERB (Equivalent Rectangular Bandwidth; Moore (1983); Glasberg and Moore (1990)) wide noise band. The noise bands were generated using Gaussian noise filtered with 4^*th*^ order gammatone filters and were separated by 1-ERB wide spectral gaps. The duration of the noise bands was 300 ms with 2 ms on-/ offset ramps (raised-cosine window). The dark bars indicate bands with fully correlated noise and interaural disparities.

**Fig. 1.**
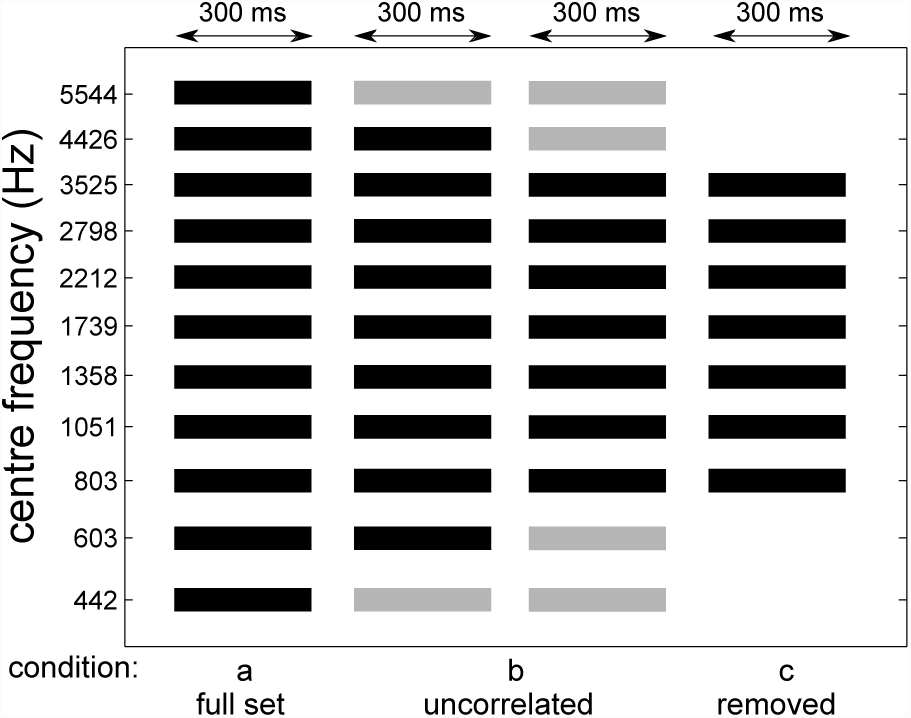
Schematic of the experimental conditions used in experiment 1. The black bars indicate noise bands with interaural disparities and the grey bars indicate bands with interaurally uncorrelated noise. All bands had a spectral width of 1-ERB and were spaced 1-ERB from each other. All bands had a duration of 300 ms.

In experiment 1a, the centre frequencies of the noise bands ranged from 442 Hz to 5544 Hz. On all bands interaural disparities were added. The ITDs ranged from −500 *µ*s to +500 *µ*s in steps of 100 *µ*s and the ILDs from −5 dB to +5 dB in steps of 1 dB, leading to 11 ITD and 11 ILD values, where negative values indicate a leading left ear and positive values a leading right ear.

The trials were generated by applying different interaural disparities to the noise bands, leading to a signal with different binaural information in the 11 frequency bands. The 11 ITD and ILD values were permuted over 11 trials and applied to the noise bands, resulting in a block of 11 stimuli with every interaural disparity occurring once per noise band. The procedure of generating blocks was repeated 11 times, leading to 121 trials per run. Each of the runs was repeated 5 times with a randomized order of the 121 trials. For each trial the same noise instances were used for all repetitions and listeners. Furthermore, the same permutations of the binaural disparities across the frequency bands were presented to the listeners but in different random orders.

In experiment 1b, the most outer or the two most outer noise bands (referred to as the “edge bands”) were presented as interaurally uncorrelated noises. Thus, these frequency bands contained no useful information for lateralization. The light grey bars in Figure 1 indicate the bands with binaurally uncorrelated noise. In experiment 1c, the two most outer bands were removed.

The stimuli were presented over headphones (Sennheiser HDA200; Sennheiser electronic GmbH & Co. KG, Wedemark, Germany) with an overall level of 60 dB SPL. The headphones were equalized with an 2048 point FIR filter modeling the inverted headphone transfer function measured on a B&K 4153 artificial ear (Brüel&Kjær Sound&Vibration Measurement A/S, Nærum, Denmark). The subjects were placed in a single-walled sound proof booth. After each stimulus they were asked to indicate if the sound was perceived as coming from the left or from the right side (1-interval, 2-alternative forced choice). A single run took the subjects about 3 to 4 minutes and breaks were allowed after every run. Sessions lasted a maximum of 2 hours.

To derive the spectral weights for the interaural disparities, a model was assumed where the physical input data (ITD and ILD values over centre frequencies) was mapped to the psychological output data (subject responses). The frequency dependent mapping variable was weight that the listener put on a particular spectral region during the decision. The mapping variable (or spectral weight) was determined by using a multiple linear regression analysis between the interaural disparities and the listeners reponse. The multiple linear regression analysis was performed using the MATLAB (The MathWorks Inc., Natick, USA) function *fitlm()* which returns the weights. These weights were normalized with respect to the average of all weights over frequency.

### 2.2 Experiment 2: Spectral weights for dynamic spectro-temporal context

In experiment 2, a paradigm inspired by auditory enhancement experiments was adopted to investigate the effect of spectro-temporal context on the spectral weights of ITD and ILD. The enhancement of a specific frequency band (referred to as on-frequency band) was promoted by using pre- and post-cursors. Figure 2 shows an example schematic of the stimulus. The procedures to create the target (black bars) and to determine the spectral weights were the same as in experiment 1. The cursors (grey bars) were diotic noise bands with a bandwidth of 1-ERB and a duration of 300 ms. Two cursors were presented before and after each off-frequency band. A gap of 2 ms was introduced between each cursor and between the cursor and the target. Five on-frequency bands were tested separately: 442, 803, 1739, 3525, and 5544 Hz.

**Fig. 2.**
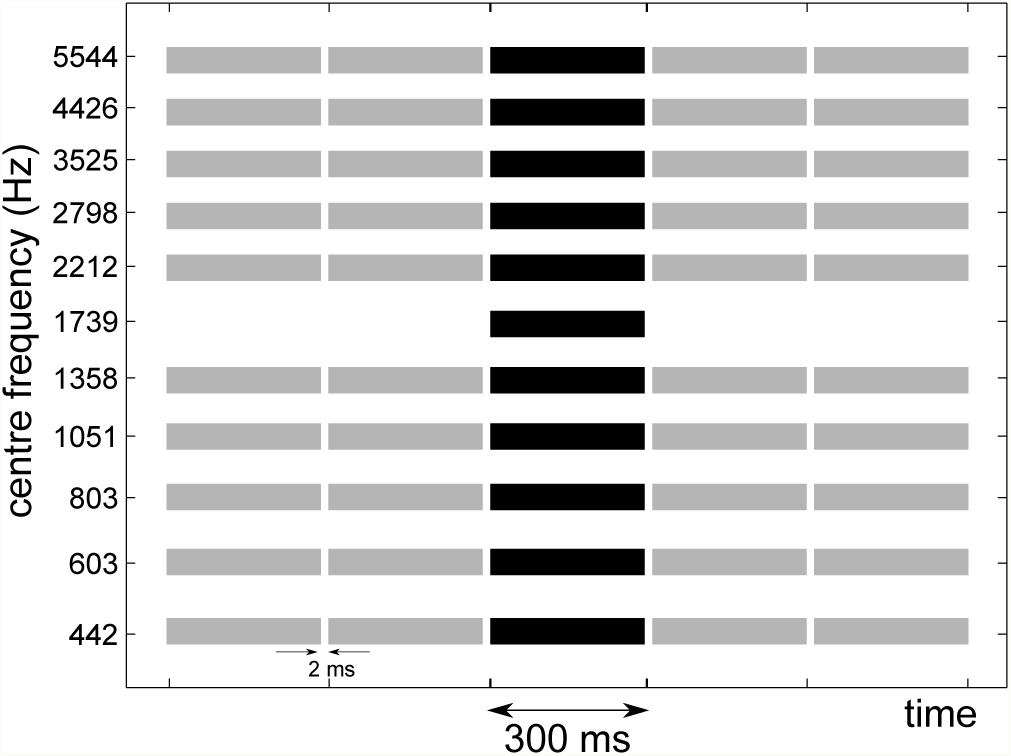
Schematic of one of the stimuli used in experiment 2. The black bars indicate noise bands with interaural disparities and the grey bars indicate the pre- and post-cursors. The cursor bands contained diotic noise. All bands had a spectral width of 1-ERB and were spaced 1-ERB from each other. All bands had a duration of 300 ms with 2 ms gaps between bands. In this example, the on-frequency band was the 1739Hz band. The remaining bands are referred to as off-frequency bands. On-frequency bands at 442, 803, 3525, and 5544 Hz were also tested.

### 2.3 Listeners

Ten listeners (6 female, 4 male) participated in experiment 1 and a subset of 6 listeners in experiment 2. The listeners were paid on an hourly basis. The hearing thresholds of the listeners measured prior to the experiment showed audiometrically normal hearing (<20 dB HL between 125 Hz and 8 kHz). The participants were between 24 and 30 years old (average 25.2 years). Four of the listeners had previous experience with psychoacoustic experiments. All participants provided informed consent and all experiments were approved by the Science-Ethics Committee for the Capital Region of Denmark (reference H-KA-04149-g).

### 2.4 Statistical analyses

The statistical analyses were performed using the statistical computing software R (version 3.6.1). Linear mixed effects models were fitted to either of the interaural disparities using the *lmerTest* package (Kuznetsova et al. 2014). The effect of the listeners was treated as a random factor. If within factor comparisons were performed, the *emmeans* package (Lenth 2016) was used with the Satterthwaite method to calculate the degrees of freedom. The post-hoc p-values were corrected for multiple comparisons using the Bonferroni correction. In the plots, asterisks and dots are used to indicate the p-values. Three asterisks indicate a p-value smaller than 0.001, two asterisks smaller than 0.01, a single asterisk smaller than 0.05 and a dot a p-value between 0.05 and 0.1. If no asterisk is plotted in the figure, no significant difference was found.

## 3 Results

### 3.1 Experiment 1: Spectral weights in static condition

#### 3.1.1 Spectral weights

Figure 3 shows the spectral weights for ITDs (panel A) and ILDs (panel B) with respect to the centre frequencies of the noise bands. The analysis of a linear mixed model showed a significant effect of the frequency bands for both ITD [*F(10,99) = 22.37, p* < *0.0001*] and ILD [*F(10,99) = 17.93, p* < *0.0001*]. The statistical values of the across frequency comparisons (post-hoc analysis) are shown in Tables 1 and 2 for the spectral weights of ITDs and ILDs, respectively. P-values in the Tables with a value *≤* 0.05 are indicated in grey.

**Table 1.**
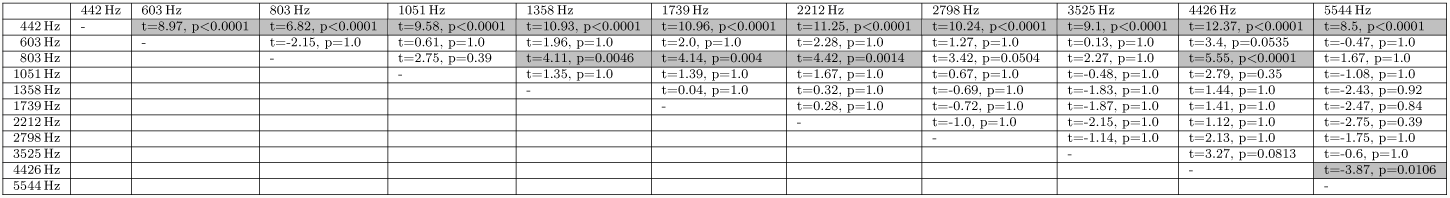
The t-ratio and the corrected p-values from the multiple comparison analysis of the spectral weights for the ITDs. The degrees of freedom are 99. Combinations with a p-value smaller 0.05 are indicated in grey.

**Table 2.**
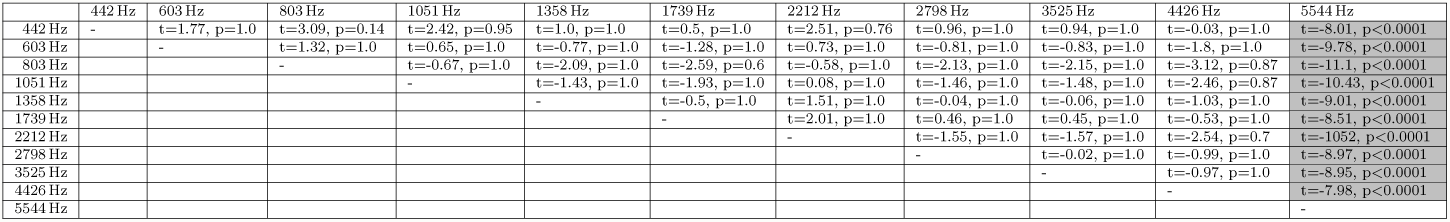
The t-ratio and the corrected p-values from the multiple comparison analysis of the spectral weights for the ILDs. The degrees of freedom are 99. Combinations with a p-value smaller 0.05 are indicated in grey.

**Fig. 3.**
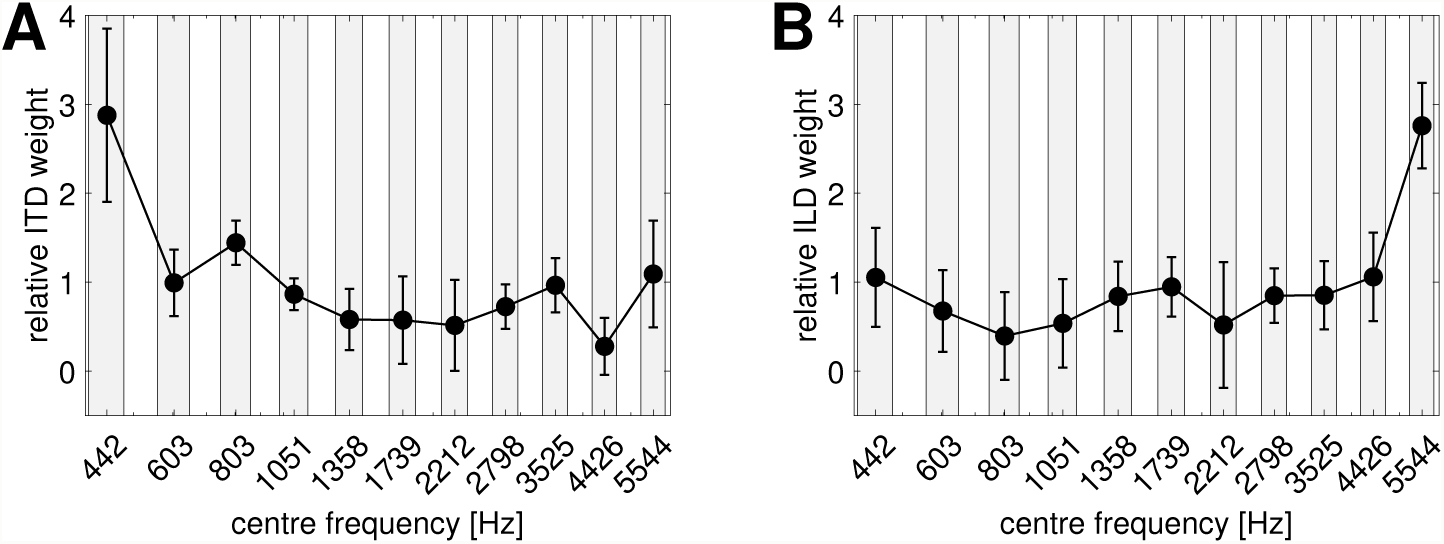
Spectral weights of ITDs (panel A) and ILDs (panel B) with respect to the centre frequencies of the 1-ERB noise bands. The circles and the errorbars indicate the mean and the standard deviation for the ten listeners, respectively. The results from the post-hoc analysis are shown in Tables 1 and 2.

The spectral weight for ITDs at the lowest frequency band (442 Hz) was significantly higher than for all other bands (see Table 1). The weight at the 803 Hz band was found above the weights for the frequency bands from 1051 to 3525 Hz as well as for the 4426 Hz band. The weight for the 4426 Hz band was also found to be lower than for the highest band (5544 Hz). The highest significant spectral weight for ILDs (see Table 2) was at the highest frequency band (5544 Hz). No other significant across-frequency difference were found for the spectral weights of ILDs.

#### 3.1.2 Effect of uncorrelated edge bands

Figure 4 shows the spectral weights for the reference conditions (experiment 1a, as in Figure 3) and the weights for experiment 1b with interaurally uncorrelated noise bands at the spectral edges. The leftwards and rightwards pointing triangles indicate the conditions with two (the highest and the lowest) and four (the two highest and the two lowest) interaurally uncorrelated noise bands as edge bands, respectively. A linear mixed model of the ITD weights revealed that the frequency [*F(10,243) = 26.65, p* < *0.0001*] and the conditions [*F(2,243) = 4.87, p = 0.0084*] were significant effects, while the interaction was not significant [F(14,243) = 0.99, p = 0.47]. A linear mixed model of the ILD weights revealed that the frequency [*F(10,243) = 11.58, p* < *0.0001*] was significant, while the conditions [*F(2,243 = 0.42, p = 0.66*] and the interaction [*F(14,243) = 1.25, p = 0.24*] were not significant. Thus, presenting uncorrelated noise bands at the edges of the spectrum did not change the contour of the weighting functions.

**Fig. 4.**
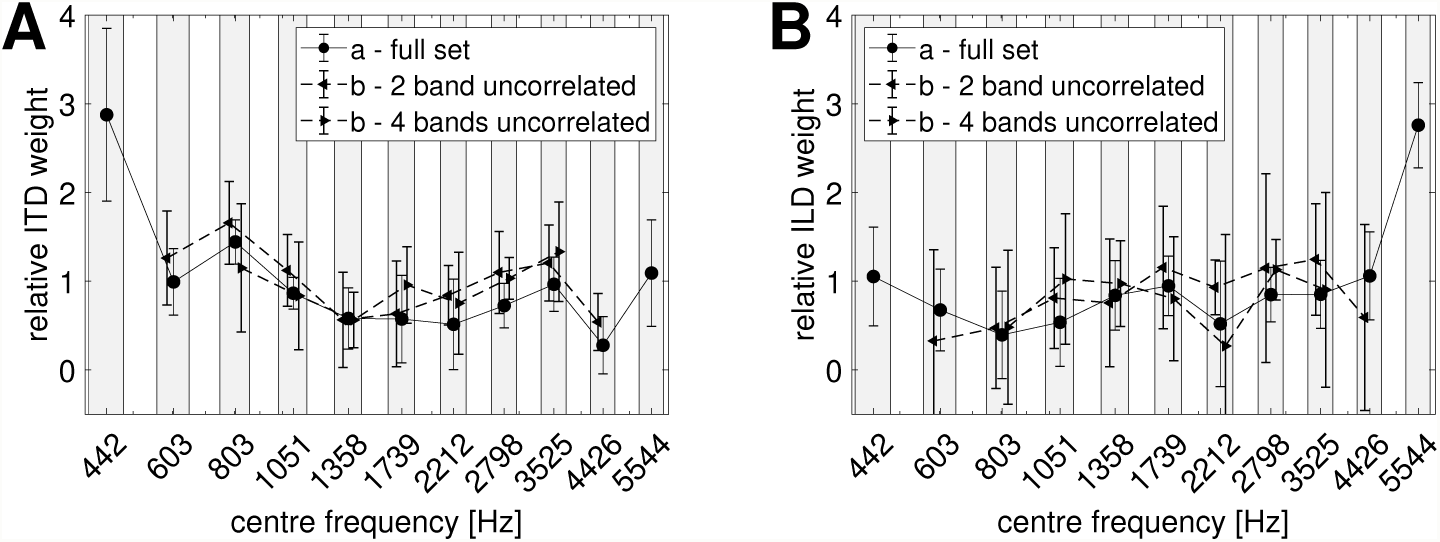
Spectral weights of ITDs (left) and ILDs (right) with respect to the centre frequencies of the 1-ERB noise bands. The circles indicate the condition with interaural disparities on all eleven bands (full set), the triangles indicate the condition with uncorrelated noise on the most outer frequency bands. The symbols and the errorbars indicate the mean and the standard deviation for the ten listeners, respectively.

#### 3.1.3 Effect of reduced stimulus bandwidth

Fig. 5 shows the spectral weights in the reference condition (experiment 1a, circle symbols) and the condition with removed edge bands (experiment 1c, diamond symbols). The linear mixed model of the ITD weights showed significant effects of both frequency [*F(10,162) = 53.63, p* < *0.0001*] and condition [*F(1,162) = 6.27, p = 0.013*] as well as of their interaction [*F(6,162) = 15.54, p* < *0.0001*]. The multiple comparison analysis of the differences between the two conditions revealed that only a significant difference at the, in this condition, lowest frequency band (803 Hz) was found [*t(162) = −9.75, p* < *0.0001*]. For the ILD weights the model showed significant effects of frequency [*F(10,162) = 26.27, p* < *0.0001*], condition [*F(1,162) = 15.31, p = 0.0001*] and their interaction [*F(6,162) = 8.51, p* < *0.0001*]. The effect of increased weights was significant both at the lowest (803 Hz) [*t(126)=-4.76, p* < *0.0001*] and at the highest (3525 Hz) frequency band for this condition [*t(126)=-6.37, p* < *0.0001*].

**Fig. 5.**
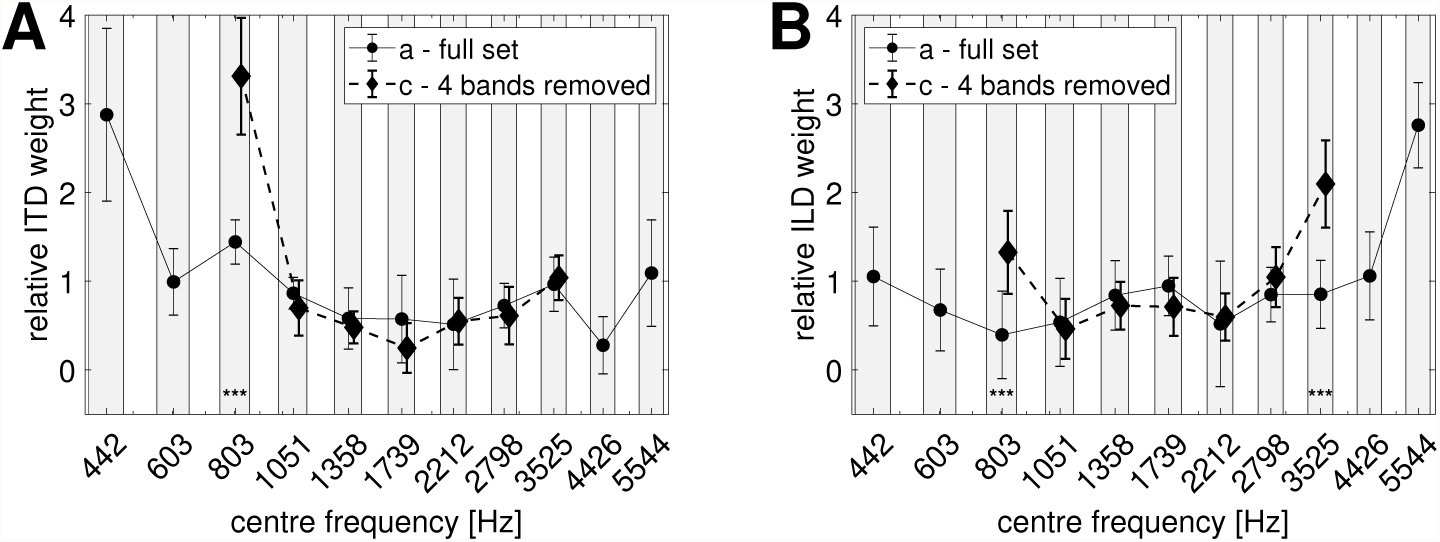
Spectral weights of ITDs (left) and ILDs (right) with respect to the centre frequencies of the 1-ERB noise bands. The circles indicate the condition with interaural disparities on all eleven bands (full set), the diamonds indicate the condition with removed stimuli on the two most outer frequency bands. The symbols and the errorbars indicate the mean and the standard deviation for the ten listeners, respectively.

### 3.2 Experiment 2: Spectral weights for dynamic spectro-temporal context

Figure 6 shows the spectral weights of ITDs (left) and ILDs (right) in the condition with pre- and postcursors. The reference condition from experiment 1a is shown as circles, the off-frequency weights are indicated in grey and the on-frequency weights in black. The symbols at the bottom of the figures indicate the significance level between the reference condition and the on-frequency weight.

**Fig. 6.**
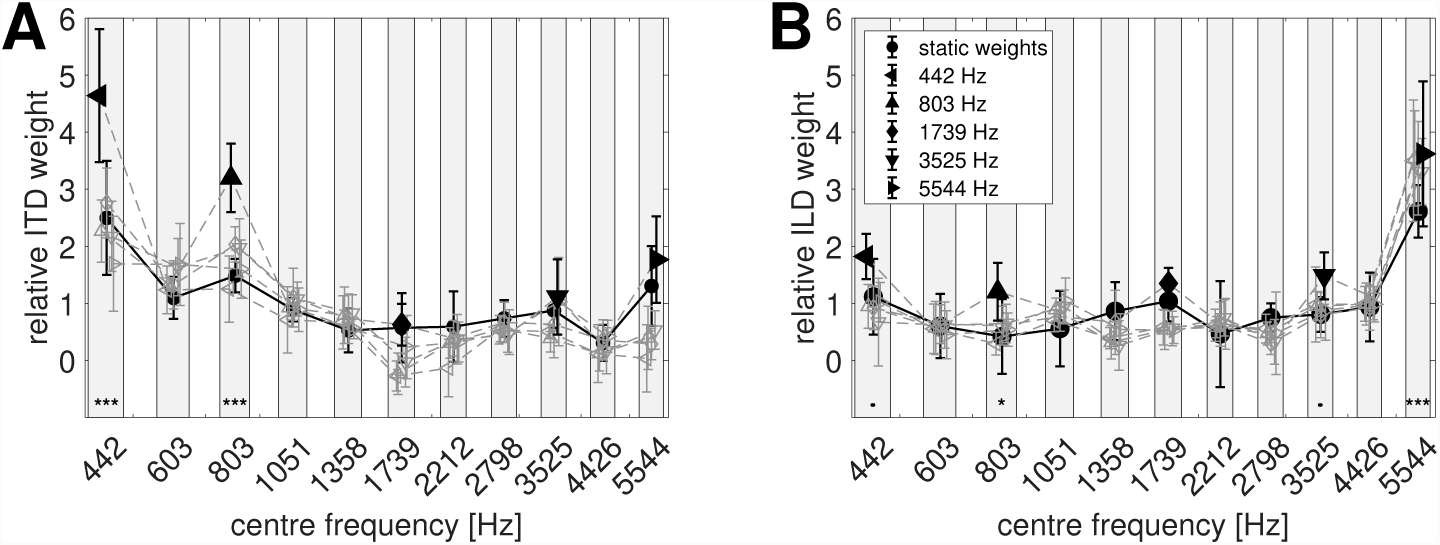
Spectral weights of ITDs (left) and ILDs (right) with respect to the centre frequencies of the 1-ERB noise bands. The circles indicate the condition with interaural disparities on all eleven bands (experiment 1a). The grey lines and symbols indicate the conditions with enhanced bands, where the black symbol represents the on-frequency band. The symbols and the errorbars indicate the mean and the standard deviation for the six listeners, respectively.

The linear mixed model of the ITD weights revealed significant effects of frequency [*F(10,325) = 89.82, p* < *0.0001*] but not of condition [*F(5,325) = 0.88, p = 0.49*]. The interaction between the two main effects was significant [*F(50,325) = 5.94, p* < *0.0001*]. The comparison of the on-frequency weights with the reference condition revealed significant increases for the 442 Hz band [*t(325)=-7.6, p* < *0.0001*] and the 803 Hz [*t(325)=-6.07, p* < *0.0001*] band but not for the 1739 Hz [*t(325)=-0.84, p = 1.0*], 3525 Hz [*t(325)=-0.18, p = 1.0*] and 5544 Hz [*t(325)=-1.64, p = 0.51*] bands.

The analysis of the linear mixed model of the ILD weights showed a significant effect of the frequencies [*F(10,330) = 100.52, p* < *0.0001*] but not of the conditions [*F(5,330) = 0.25, p = 0.94*]. The interaction between the frequencies and the conditions was significant [*F(50,330) = 1.79, p = 0.0016*]. Comparing the on-frequency weights to the corresponding weights in the reference condition showed a significant increase of 0.86 of the weight at the highest tested frequency band [5544 Hz: *t(330)=-3.62, p = 0.0017*]. The 3525 Hz bands showed an average increase of 0.63, however the p-value was found above the significance level of 0.05 [*t(330)=-2.43, p = 0.0782*]. The weights at the 1739 Hz band were not significantly affected by the enhancement [*t(330)=-1.11, p = 1.0*]. The weights at 803 Hz [*t(330)=-2.83, p = 0.0247*] and 442 Hz [*t(330)=-2.54, p = 0.0578*] increased on average by 0.81 and 0.77.

## 4 Discussion

In the present study the spectral weighting for a stimulus consisting of 11 simultaneously presented 1-ERB wide noise bands with ITDs and ILDs was investigated. It was shown that the highest weight for ITDs was given to the frequency band with the lowest centre frequency, and at the highest weight for ILD was given to the frequency band with the highest centre frequency. The remaining bands received substantially lower weights than these edge bands. This “edge effect” was also found when reducing the overall bandwidth of the stimulus. When presenting interaurally uncorrelated noise as the edge bands, no change in weight was observed, resulting in a weighting function with equal spectral weights. The auditory enhancement paradigm in experiment 2 led to an increase of the on-frequency band. This enhancement of the weight was found for ITDs at low frequencies and for ILDs at low and high frequencies.

The results from this study are in general agreement with the Duplex Theory (Rayleigh 1907; Macpherson and Middlebrooks 2002): ITDs receive the highest weight at low frequencies and ILDs at high frequencies. However, when assuming that ITD information is prominent at low frequencies and that the amount of useful information gradually decreases towards high frequencies (Klumpp and Eady 1956; Brughera et al. 2013), and vice versa for ILDs (Mills 1960), the findings of the present study show a different pattern. Instead of a gradual change of the weights, a sharp transition from high to low weights was found at the frequency bands located at the spectral edges of the stimulus. This is at odds with estimates of spectral weights based on ITD and ILD thresholds where a gradual change of the spectral weight for ITDs has been shown (Raatgever 1980; Stern et al. 1988; Buchholz et al. 2018). A reason for this difference might be that the listeners in the current study had to integrate the binaural information across frequencies and, thus, binaural interference (McFadden and Pasanen 1976) was taken into consideration.

The current study was designed to reduce within-channel interference by separating the 1-ERB wide noise bands by 1-ERB wide spectral gaps. Thus, only little energy “leaked” into neighbouring auditory channels. However, binaural auditory filters have been shown to be wider than monaural auditory filters (van de Par and Kohlrausch 1999; Kolarik and Culling 2010; van der Heijden and Trahiotis 1998; Bernstein and Oxenham 2006; Holube et al. 1998). Assuming wider binaural auditory filters, this might have introduced a within-channel interference of the interaural disparities. This possible interference might have led to a reduction of the weight in a given frequency band as conflicting binaural information from two or more bands were integrated within one binaural auditory filter. Thus, the frequency bands at the spectral edges with only a single neighbouring band were less affected by within-channel interference than the remaining bands. This explanation is supported by the conditions with altered edge frequency bands where band were removed or replaced with uncorrelated noise. When a band was removed, the interference for the new edge band is reduced and thus the spectral weight of this band increases. However, when there is uncorrelated noise on the edge band, the interference remains constant and thus the weight is unchanged. However, the edge effect only occurred if usable interaural information for the auditory system were available. This was the case mainly in low frequencies for ITDs and in high frequencies for ILDs. Yet, in the condition with removed edge bands, also the lowest frequency band for ILDs was increased in comparison to the reference condition, which suggests that enough ILD information was available at the mid frequency range but not at the low frequency range.

In experiment 2, spectral weights were determined while using an auditory enhancement paradigm (Viemeister 1980; Byrne et al. 2011). The weights at the cued bands were found to be increased at low and high frequencies for ILDs and at low frequencies for ITDs with respect to the reference condition. The results are in line with findings on ITD and ILD detection thresholds as discussed above, but not with the the duplex theory as ILD weights are increased also at low frequencies.

Comparing the weights across the enhanced bands, it is apparent that the edge frequency bands at low and high frequencies receive the highest weights for ITDs and ILDs, respectively. This is not in line with threshold measurements of ITDs and ILDs. While ILD thresholds are constant over frequency (Yost 1988; Grantham 1984; Rowland and Tobias 1967), ITD thresholds have been shown to be lowest at 800 Hz (Klumpp and Eady 1956; Brughera et al. 2013).

The reason for the change of the spectral weights using the auditory enhancement paradigm remains unclear. One possibility could be due to an internal gain of the enhanced band, leading to louder perceived band (Viemeister 1980) and therefore to a higher weight. Second, both the enhancement and the binaural processing might happen at the same stage for example in the inferior colliculus (Nelson and Young 2010), which might lead to a more efficient coding of information in the enhanced band. Third, the reason for increased weights could be due to a reduced binaural interference. Best et al. (2007); Woods and Colburn (1992) showed that binaural interference is reduced when auditory information is grouped. However, grouping has been argued to not be a reason for auditory enhancement (Summerfield et al. 1987; Byrne et al. 2011), thus, it might also not be the underlying mechanism for the increased spectral weights. The current study can neither prove nor rule out any of the three reasons and further work is needed to link auditory enhancement and binaural perception.

In the current study the spectral weighting of ITDs and ILDs has been investigated in separation. However, to lateralize a sound, both ITDs and ILDs are used jointly. Thus, one might investigate a common lateralization weighting function. Wightman and Kistler (1992) showed that low-frequency ITDs are dominant for sound source localization. If the auditory system also weights low-frequency ITD information higher than other localization cues remains unknown. Generally, we think that the analysis of separate weighting functions for ITDs and ILDs is valuable as they have been shown to be processed in different nuclei along the auditory pathway (Yin 2002).

Even though, the edge frequency bands have been found to receive the highest weight, also the other weights were found to be above zero. Thus, all frequencies were found to contribute to the lateralization.

## 5 Conclusions

In the current study we investigated the across-frequency integration of interaural time- and level differences. An observer-weighting analysis paradigm was used where spectral weights of ITDs and ILDs were determined using a multiple linear regression approach. It has been shown that ITD weights are largest at the lowest frequency band and ILD weights are largest at the highest frequency band. When using an auditory enhancement paradigm these weights increase at the enhanced frequency band while the remaining frequencies remain constant. However, the change in the weighting function was only observed at low frequencies for ITDs but for both low and high frequencies for ILDs.

